# Multi-hops functional connectivity improves individual prediction of fusiform face activation via a graph neural network

**DOI:** 10.1101/2020.09.21.305839

**Authors:** Dongya Wu, Xin Li, Jun Feng

## Abstract

Brain connectivity plays an important role in determining the brain region’s function. Previous researchers proposed that the brain region’s function is characterized by that region’s input and output connectivity profiles. Following this proposal, numerous studies have investigated the relationship between connectivity and function. However, based on a preliminary analysis, this proposal is deficient in explaining individual differences in the brain region’s function. To overcome this problem, we proposed that a brain region’s function is characterized by that region’s multi-hops connectivity profile. To test this proposal, we used multi-hops functional connectivity to predict the individual face response of the right fusiform face area (rFFA) via a multi-layers graph neural network and showed that the prediction performance is essentially improved. Results also indicated that the 2-layers graph neural network is the best in characterizing rFFA’s face response and revealed a hierarchical network for the face processing of rFFA.

## Introduction

Brain connectivity acts as the pathway for transferring information between brain regions and determines the information inflow and outflow of each cortical region. Passingham et al. (2002) proposed that the function of each cortical region can be determined by the region’s input and output connectivity profiles. Mars et al. (2018) further tested and extended this proposal via the neuroimaging of connectivity, and showed that the connectivity space composed by each region’s connectivity profiles provides a powerful framework in describing a brain region’s function.

The connectivity profile can be defined in terms of the white matter pathway represented by tractography through diffusion magnetic resonance imaging (MRI), or in terms of the temporal coupling between spontaneous fluctuations of resting-state functional MRI (rfMRI) signal. Under the proposal of Passingham et al. (2002), previous studies have utilized structural connectivity (Johansen-Berg et al. 2004; Tomassini et al. 2007; Beckmann et al. 2009; Saygin et al. 2011a) or functional connectivity (Cohen et al. 2008; Gordon et al. 2016) to characterize the boundary of functionally distinct brain regions, or have utilized structural connectivity (Saygin et al. 2011b; Osher et al. 2016; Saygin et al. 2016; Wu et al. 2020) or functional connectivity (Tavor et al. 2016; Parker Jones et al. 2017) to predict the functional activation information of brain regions at various task states.

Though the proposal that a brain region’s function is represented by the input and output connectivity profiles is widely adopted in various studies, this proposal is deficient in characterizing the individual brain region’s function. Specifically, under this proposal, a brain region’s function can be represented by a linear combination of the region’s connectivity profiles, with the weight of the connectivity profiles viewed as the functional information transferred from neighboring regions. However, the weight of the linear model is the same for all subjects. This amounts to assume that the functional information of neighboring regions is the same for different subjects. In other words, the prediction model under this proposal neglects individual differences in the functional information of neighboring regions and thus is deficient in predicting individual brain region’s function.

To overcome this problem, we proposed that more connectivity features in the brain connectivity network should be considered. Explicitly, as shown in Fig. 1A, the functional activation information of 1-hop regions is unknown, however, the information of 1-hop regions is transferred from 2-hop regions through the connections of 1-hop regions. According to this logic, when the functional activation information of all brain regions is unknown, the connections of n-hop regions may contain functional information of the region of interest (ROI). Denote the direct input and output connections of the ROI as the 1-hop connections. The connections of the 1-hop regions amount to the 2-hop connections of the ROI. We then define the ensemble of these 1-hop and 2-hop connections as the 2-hops connections relative to the ROI. Using these terms, the previous proposal (Passingham et al. 2002) is formulated as that a brain region’s function is represented by the 1-hop connectivity profiles. Separately, based on the above analyses, we proposed that a brain region’s function is represented by the multi-hops connectivity profiles.

**Figure 1.**
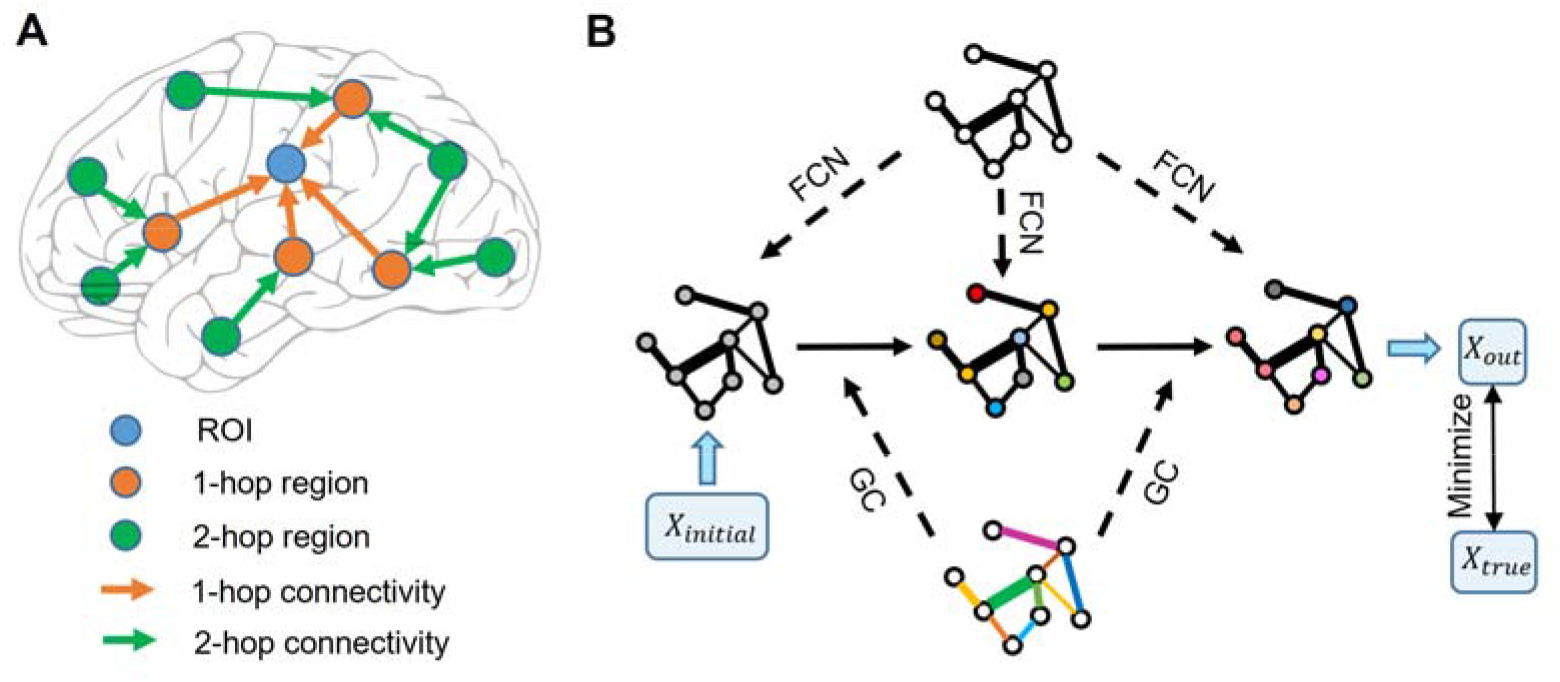
Schematic illustration of graph neural network. (A) Functional information of the ROI is transferred from the 1-hop neighboring regions via the 1-hop connections, and the functional information of the 1-hop neighboring regions is transferred from the 2-hop neighboring regions via the 2-hop connections. (B) Node color represents the functional information of brain regions. Edge width represents the strength of functional connectivity pathways. Edge color represents the coefficient of graph convolution (GC), i.e. the extent to which each connection participates in the functional information propagation. Initial functional information transfers within the functional connectivity network (FCN) through the graph convolution network. Coefficients of GC are trained by minimizing the error between the predicted output and true functional activation information.

To further test our proposal, we selected the right fusiform face area (rFFA) as the ROI. We adopted the FACES-SHAPES (emotion task) and FACE-AVG (working memory task) contrasts in the human connectome project (HCP) to define individual subject’s functional face selectivity, and utilized the rfMRI data to construct individual brain functional connectivity network. Inspired by the fact that computations in the graph neural network are analogous to the propagation of functional information in the brain connectivity network, we designed a multi-layers graph neural network (Fig. 1B). This graph neural network is well suitable for our proposal because it includes both direct and indirect, single-step and multiple-step connectivity features to characterize functional selectivity of the ROI. Finally, we applied the graph neural network containing the multi-hops functional connectivity to predict individual face selectivity of the rFFA.

## Materials and Methods

### Human Connectome Project data

We used the minimally pre-processed data (Glasser et al. 2013) provided by the HCP S1200 release. We selected all the 997 subjects that have the FACES-SHAPES, FACE-AVG task contrasts, and resting-state fMRI acquisitions.

Task and resting-state fMRI data were projected onto 2 mm standard CIFTI grayordinates space, and the multimodal surface matching (MSM) algorithm (Robinson et al. 2014) based on areal features (MSMAll) was used for accurate inter-subject registration. Acquisition parameters and processing are described in detail in several publications (Barch et al. 2013; Smith et al. 2013). Briefly, resting and task fMRI scans were acquired at 2 mm isotropic resolution, with a fast TR sampling rate at 0.72 s using multiband pulse sequences (Ugurbil et al. 2013). Both sets of functional data had already been registered to the MNI space (Glasser et al. 2013). Each subject had four 15-minute resting fMRI runs, with a total of 1,200 time points per run. The resting fMRI data were further pre-processed by FIX to automatically remove the effect of structured artifacts (Griffanti et al. 2014; Salimi-Khorshidi et al. 2014).

### Functional Connectivity profile

We calculated functional connectivity based on the HCP-MMP1.0 (Human Connectome Project Multi-Modal Parcellation version 1.0) (Glasser et al. 2016) that containing 360 brain regions. The functional connectivity was calculated from the resting fMRI data. The four runs of individual resting-state time series data were concatenated after being demeaned and variance-normalized along the time axis. We did not apply global signal regression before calculating functional connectivity. The time series for every vertex in the seed region was correlated with the averaged time series for each of the remaining 359 target regions. The diagonal elements of the functional connectivity matrix were set as ones.

### Gaussian-Gamma mixture model for determining activation threshold

The HCP_MMP1.0 contains 360 brain regions. Using a connectivity matrix with size 360×360 is likely to overfit the training dataset. Since some brain regions do not involve in the face recognition process, removing these brain regions beforehand can reduce the model complexity in a great deal. To prevent some individual differences with opposite signs from canceling with each other, we calculated each brain region’s mean absolute activation across subjects. Then we used the Gaussian-Gamma mixture distribution (Gorgolewski et al. 2012) to model the density distribution of the mean absolute activation for the 360 brain regions. The Gaussian distribution is used to model the random noise effect, and the Gamma distribution is used to model the activation signal. We set the activation threshold as the point when the probability density of Gamma distribution is higher than that of Gaussian distribution.

### Graph neural network for predicting functional face selectivity

Graph neural network is widely used to process data with graph structures (Defferrard et al. 2016; Kipf and Welling 2017; Veličković et al. 2018). We developed a graph neural network that is adapted to process brain connectivity network data (Fig. 1B). The graph neural network with a single-layer can be represented in a matrix form as follows:

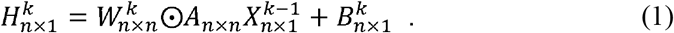

*k* represents the *k*-th layer of the network. *n* represents the number of nodes in the graph neural network, i.e. the number of brain regions. *X* and *H* respectively represent the input and output of the graph neural network, i.e. the functional activation of each brain region. *B* represents the bias term of the model. *A* represents the adjacency matrix composed of functional connectivity. *W* represents the functional information propagation coefficient of each functional connectivity pathways. 147 exerts on *A* via the operation ⊙ that represents the element-wise multiplication. A multi-layers computation can be achieved by applying formula (1) repetitively. When one trains a multi-layers graph neural network, the vanishing gradient effect usually occurs. Inspired by the residue neural network (He et al. 2016), we added residue connections between neighboring layers to formula (1) and the resulting model is:

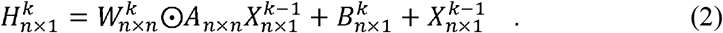

To make the 1-layer graph neural network model consistent with the linear model adopted by previous studies, the brain region’s activation in the 0th layer is initialized as all ones: 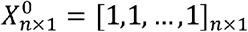. It is worth noting that model (2) doesn’t involve nonlinear activation functions but is nonlinear in the sense that the functional connectivity feature *A* occurs in every computation layer. Therefore the resulting multi-layers model contains nonlinear features that represent functional information passing through multi-steps functional connectivity pathways.

### Metrics for assessing the model

We used two metrics to assess the model performance on testing data. Denote the target value by *y* and the prediction value by 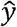. The sum squared error 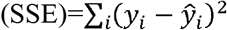 is widely used to assess the difference between the target and prediction values. But in different divisions of the dataset, the variance of the target value is different, thus the SSE cannot be compared across different divisions. We divided the SSE by the sum squared total 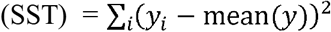 and the resulted normalized squared error (NSE) was used. We also adopted the Pearson correlation to assess the similarity between the actual and prediction value, denoted by *r*. Under the least squares condition, NSE represents the error proportion that cannot be explained by the model, and *r*^2^ represents the proportion of target data that can be explained by the model. Since the least squares condition is not satisfied by our model, these two metrics only serve as approximations.

Prediction similarity assessed by the Pearson correlation coefficient was Fisher’s *z* transformed when used for further statistical tests. Since the evaluation metrics for different models were paired for each random division of the dataset, we performed paired-sample *t*-tests using the custom Matlab command “ttest”.

### Implementation details

The whole dataset was randomly divided into a training set and a testing set with a ratio of nine to one. Though the sample size 997 is relatively large in neuroimaging, it is rather small compared to that of computer vision datasets in machine learning that usually contain over ten thousand samples. The random splitting of a small dataset can introduce random effects into the final results, i.e. the metrics for assessing the model can vary widely across different divisions. To remove the random effect as much as possible, we performed the prediction process 100 times with different random divisions of the dataset and used the mean of the two metrics to assess the model.

We implemented the graph neural network with PyTorch (https://pytorch.org/). The parameters of the model were initialized by the Xavier normal distribution with a gain of 0.1. The model was trained via the stochastic gradient descent optimizer with a Nesterov momentum of 0.9 used. The training batch size was 128 and 500 training epochs were used. The initial learning rate was 0.01 and a 0.1 multiplicative factor of learning rate decay was set at 300 and 400 epochs respectively. To further overcome the problem of overfitting caused by a small sample size, we added Gaussian random noise to the individual connectivity features at each training step. The variance of the Gaussian random noise was set equal to the variance of each connectivity feature across subjects. This technique can be viewed as a kind of online data augmentation. The graph neural network models were trained on an NVIDIA GeForce GTX 1080 Ti graphic processing unit. The training time for each random separation lasts about 8 minutes, and the total training time for 100 random separations lasts about 13 hours.

### Data availability

The HCP S1200 data release is publicly available online at https://www.humanconnectome.org/.

## Results

### Functional face ROI localization

We selected the rFFA as the ROI, given that this region is the most selective in the face processing network (Kanwisher et al. 1997). It is advantageous to study individual differences by choosing a region that has a specialized function and is reliably replicated across studies and participants. We identified the rFFA by the right fusiform face complex (rFFC) region in the HCP-MMP1.0 (Glasser et al. 2016). The FACES-SHAPES task contrast for the HCP emotion paradigm and the FACE-AVG task contrast for the HCP working memory paradigm were separately used to identify the functional face response. We showed the group average *z*-statistics of these two task contrasts and the boundary of rFFC region in Fig. 2A. The rFFC region was identified in both task contrasts, and the boundary of rFFC region coincided well with that of the group average activation. The mean *z*-statistic within the rFFC region was used to assess each subject’s face selectivity.

**Figure 2.**
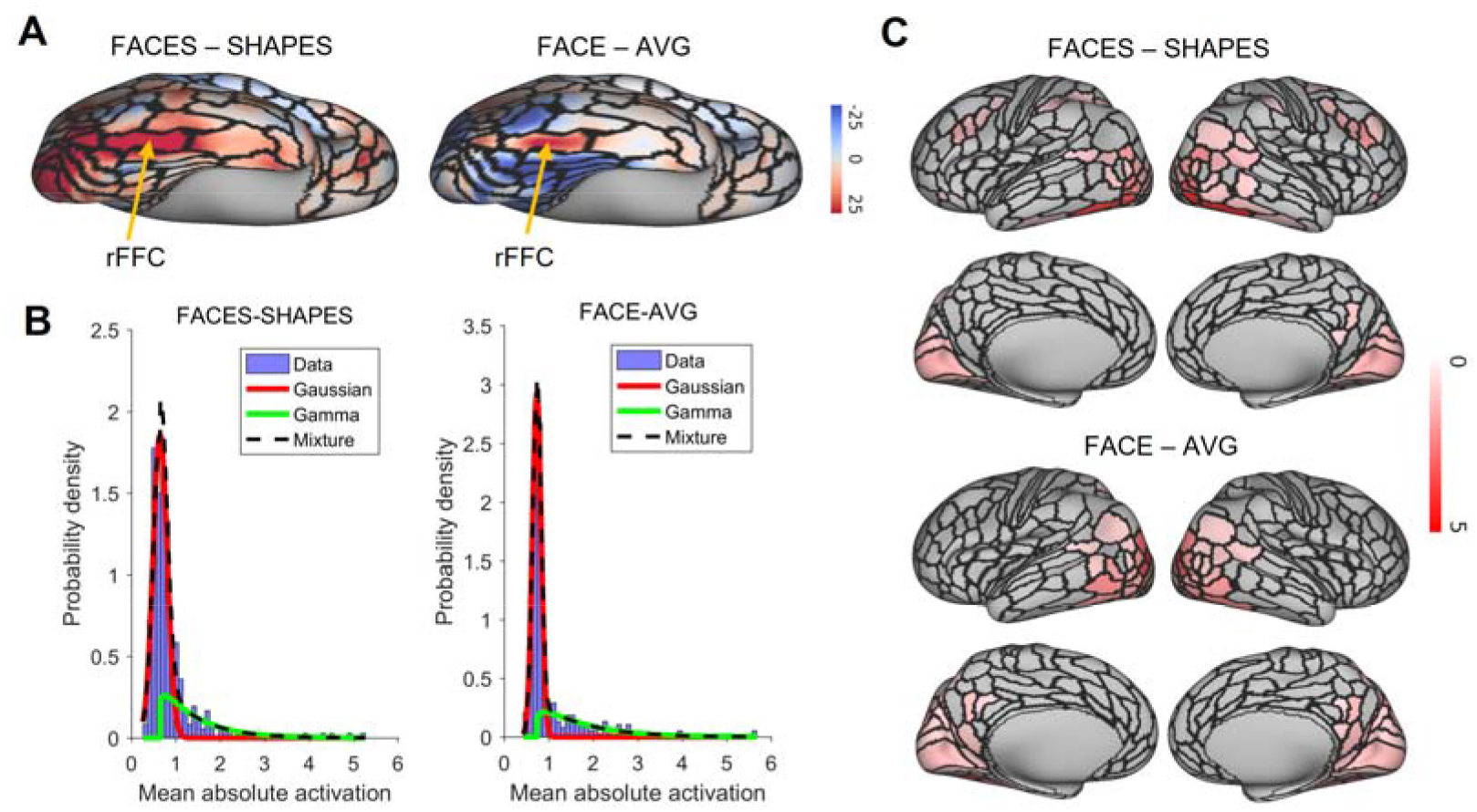
Functional face ROI localization and network selection. (A)Voxel-wise group average *z*-statistics of both contrasts were shown. The yellow arrow indicates the area where the rFFC region locates. (B) The purple histogram indicates the density distribution of the mean absolute activation for the 360 brain regions. A Gaussian (red curve)-Gamma (green curve) mixture (black curve) model was used to fit the data. (C) Mean absolute *z*-statistics of brain regions in the functional face activation network were shown.

### Functional face activation network selection

We next defined the functional face activation network in preparation for constructing the graph neural network. Fig. 2B shows the density distribution of the mean absolute activation for the 360 brain regions. We used a Gaussian-Gamma mixture model to fit the density distribution. The threshold of the activation network was chosen when the probability of Gamma distribution surpassed that of Gaussian distribution. We showed the activation networks of both task contrasts in Fig. 2C. Networks of both task contrasts mainly include brain regions in the visual cortices, such as the primary and early visual cortices, dorsal and ventral stream visual cortices, MT+ complex and neighboring visual areas. Both networks also include medial and lateral temporal cortices, superior and inferior parietal cortices, temporo-parieto-occipital junction, and posterior cingulate. In addition, the activation network of FACES-SHAPES contrast also includes inferior frontal, orbital and polar frontal, dorsolateral prefrontal, and premotor cortices. The activation network of FACES-SHAPES contrast is broader than that of FACE-AVG contrast, because the activation network of FACE-AVG contrast is mainly for basic face perception, while the activation network of FACES-SHAPES contrast also includes emotional processing of faces.

### Statistical validation of graph neural network prediction model

After selecting the functional face activation network, we constructed graph neural networks to predict individual face selectivity of the rFFC region. We first compared the 2-layers graph neural network with the random permutation model to validate the prediction model statistically. The random permutation model was constructed with the same structure as the 2-layers graph neural network, except that the pairings between the functional connectivity network and the response of rFFC region was shuffled. We illustrated the comparison between the 2-layers graph neural network and the random permutation model in Fig. 3A. For the FACES-SHAPES contrast, the prediction error of the 2-layers graph neural network is significantly (*t*(99) = -26.5, *p* = 9.5×10^-47^, paired-sample *t*-test) lower than that of the random permutation model, and the prediction similarity of the 2-layers graph neural network is significantly (*t*(99) = 33.2, *p* = 1.7×10^-55^, paired-sample *t*-test) higher than that of the random permutation model. Results for the FACE-AVG contrast are similar (see Supplementary Table S1). Hence, the 2-layers graph neural network can capture the association between the individual functional connectivity network and the face selectivity of rFFC region above random level.

**Figure 3.**
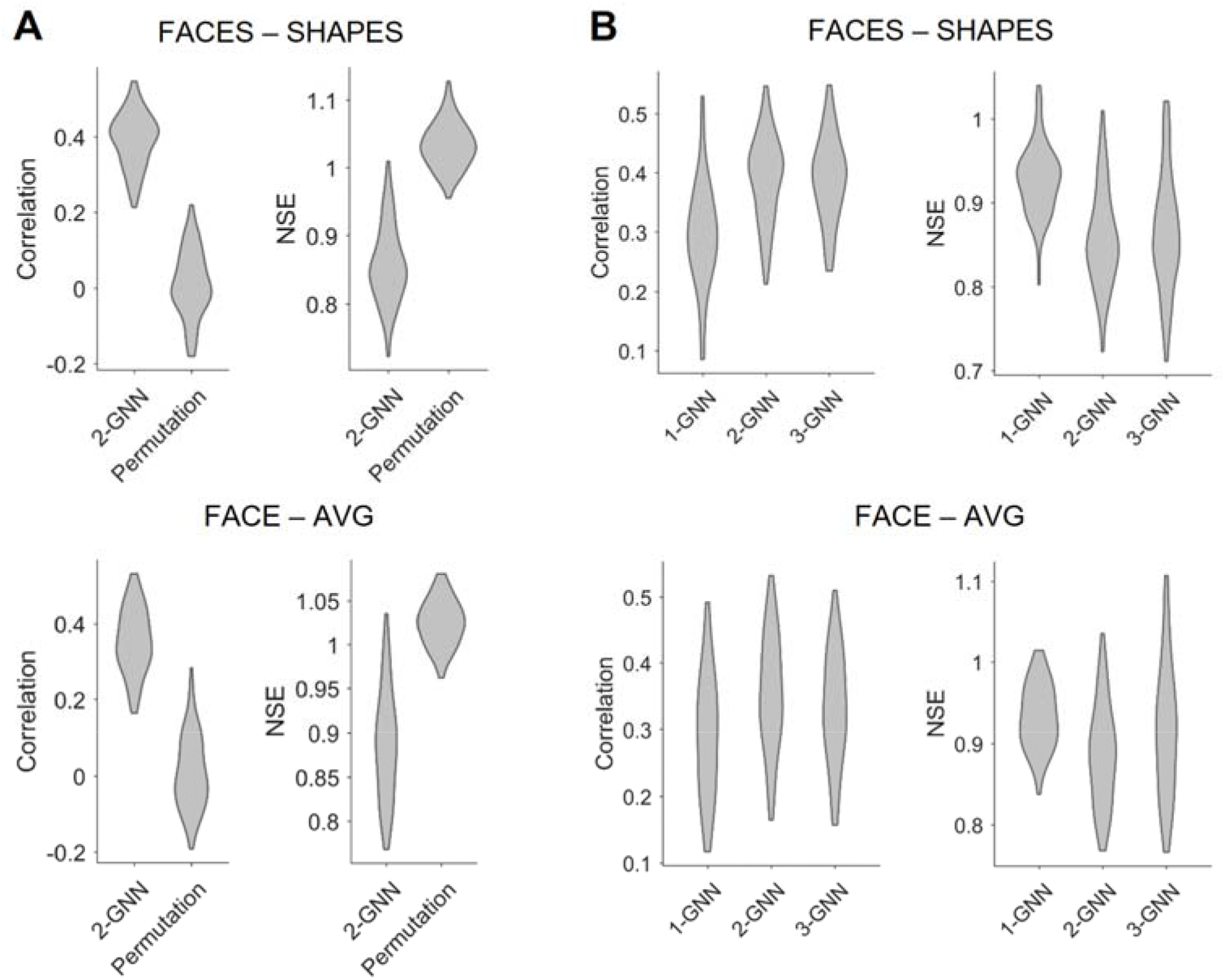
Comparison of prediction metrics for different models. (A) Comparison between the 2-layers graph neural network (2-GNN) and the random permutation model with the same structure. The prediction performance of 2-GNN is better than that of random permutation model in that the 2-GNN has a higher prediction similarity (assessed by correlation) and a lower prediction error (assessed by NSE). (B) Comparison of graph neural networks with different layers. The 2-GNN has a higher ability to predict better individual differences than both the 1-GNN and 3-GNN.

### Comparison of graph neural networks with different layers

After validating the graph neural network statistically, we further tested our proposed assumption by comparing graph neural networks with different layers. The 1-layer graph neural network corresponds to the linear prediction model adopted by previous studies and utilizes the 1-hop (i.e., direct) functional connectivity of the rFFC region to predict the rFFC region’s individual functional face response. In the 2-layers graph neural network, the final layer corresponds to the 1-hop functional connectivity representation of the rFFC region’s face response, and the first layer corresponds to the 2-hop functional connectivity representation of the rFFC region’s face response. Thus, the multi-layers graph neural network contains the representation of the rFFC region’s face response through multi-hops functional connectivity. If the proposal that a brain region’s function is represented by the multi-hops connectivity is rational, using multi-hops functional connectivity should improve the prediction of the rFFC region’s face response. We determined the rational number of hops based on the generalization ability of the graph neural networks with different number of layers and showed the comparison results in Fig. 3B and Supplementary Table S1. For the FACES-SHAPES contrast, the prediction error of the 2-layers graph neural network (mean NSE = 0.857) is significantly (*t*(99) = -16.0, *p* = 3.0×10^-29^, paired-sample *t*-test) lower than that of the 1-layer graph neural network (mean NSE = 0.927), but is not very significantly (*t*(99) = -2.09, *p* = 0.039, paired-sample *t*-test) lower than that of the 3-layers graph neural network (mean NSE = 0.864). The prediction similarity of the 2-layers graph neural network (mean correlation = 0.392) is significantly (*t*(99) = 13.5, *p* = 2.8×10^-24^, paired-sample *t*-test) higher than that of the 1-layer graph neural network (mean correlation = 0.296), but is not significantly (*t*(99) = 0.269, *p* = 0.788, paired-sample *t*-test) higher than that of the 3-layers graph neural network (mean correlation = 0.391). Results for the FACE-AVG contrast are similar, except that the differences between the 2-layers and 3-layers graph neural networks are significant (see Supplementary Table S1). Since the evaluation metrics stopped improving, we only tested graph neural networks with the number of layers up to 3. Overall, the multi-layers graph neural network improves the prediction performance in individual face selectivity of the rFFC region, and the 2-layers graph neural network possessed the best prediction performance.

### Functional network pathways involving face selectivity

In the previous section, we determined that the 2-layers graph neural network has the best generalization ability, thus the functional network pathways containing the rFFC region’s 1-hop and 2-hop functional connectivity best characterize the individual functional face selectivity of the rFFC region. Fig. 4 depicts the 2-hop functional connectivity network involving face selectivity. The 2-hop functional connectivity network also contains information about the 1-hop functional connectivity, because the output of the first layer is subsequently used as the input of the second layer. The 2-hop functional connectivity network is not symmetrical. The rows represent brain regions that integrate functional information from neighboring regions, and the columns represent brain regions that send functional information out. In both task contrasts, connectivity coefficients with large absolute values mainly concentrate in the rows of ventral stream visual cortices and MT+ complex visual areas. This result indicates that brain regions in these two cortices mainly participate in the computation of the following layer. Though some brain regions in other rows do not have large absolute connectivity coefficients, these regions in the columns have large absolute connectivity coefficients. This result indicates that some brain regions do not integrate functional information for the following layer, but they send functional information out to the regions that integrate functional information. The whole results suggest a hierarchical functional face processing mechanism for the rFFC region. The rFFC region first mainly integrates functional information from regions in the ventral stream visual cortices and MT+ complex visual areas, then regions in these two cortices integrate functional information from regions in other cortices.

**Figure 4.**
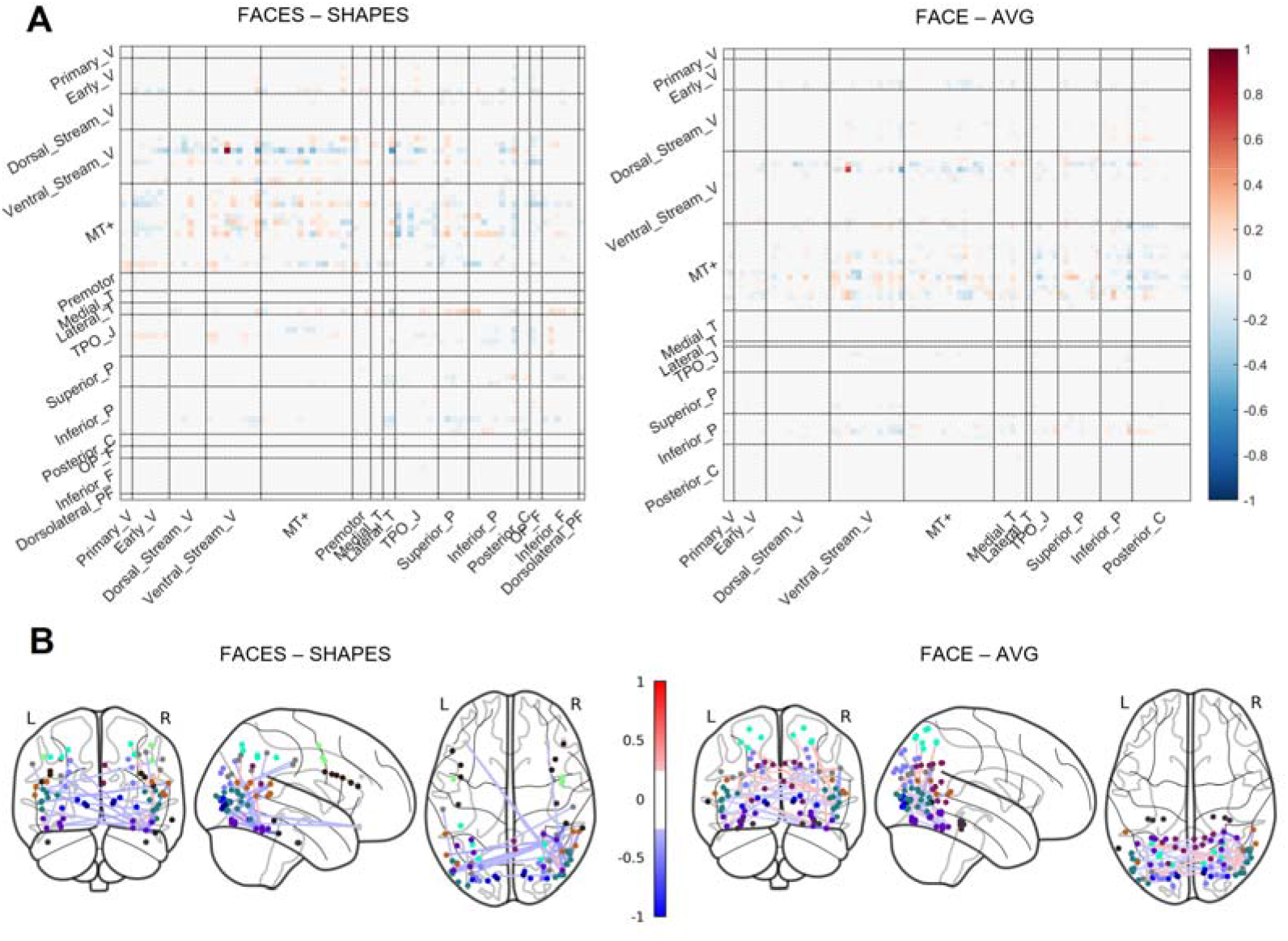
Visualization of connectivity pathways involving face selectivity. (A) The functional network involving face selectivity is shown in a matrix plot. Color represents the coefficient strength of each connection. Brain regions that belong to the same cortex are grouped. Full names of cortices are included in Supplementary Table S2. (B) The functional network is plotted on the glass brain. Only the connection with the top 1% coefficient strength is shown to make the plot clean. Edge color represents the coefficient strength of each connection. Node color represents the cortices that each brain region belongs to (see Supplementary Fig. S1).

## Discussion

In order to better characterize the brain region’s function of individuals, our study proposed that a brain region’s function is represented by the multi-hops connectivity profiles. The multi-layers graph neural network model was used to incorporate multi-hops connectivity features in the functional connectivity network. We tested our proposal by predicting the functional face response of the rFFC region via the rFFC region’s multi-hops functional connectivity. Our results showed that the 2-hops functional connectivity profile has the best generalization ability in characterizing the rFFC region’s individual functional face response, and revealed a hierarchical network for the rFFC region’s functional face processing mechanism. The current study provides new insights into understanding the brain region’s function from a network perspective.

Previous researchers proposed that a brain region’s function is represented by the 1-hop connectivity profiles (Passingham et al. 2002). However, this proposal neglects individual differences in the functional information of ROIs’ neighboring regions. Under our proposal that a brain region’s function is represented by the multi-hops connectivity profiles, individual differences in the 1-hop brain region’s functional information are taken into consideration via the 2-hop connectivity profiles. Our proposal is also consistent with neuroscience findings. Researchers have suggested that brain functions do not rely on the independent operation of a single brain region or connectivity pathway, but derive from the brain network composed of multiple brain regions and connectivity pathways (McIntosh 2000; Misic and Sporns 2016). In addition, indirect connectivity features among other brain regions can also affect ROIs via the brain network (Honey et al. 2009). Brain navigation efficiency is also due to multi-hop brain connectivity pathways (Seguin et al. 2018). In a word, since the multi-hop connectivity encodes the topological and geometrical properties of the brain connectivity network, our proposal indicates that a brain region’s function is encoded in the topology of the brain connectivity network.

The multi-layers graph neural network perfectly matches our proposal, since the multi-layers convolution computations characterize the propagation of functional information among the brain connectivity network. Though some kinds of graph neural network models have been developed to process brain network data (Ktena et al. 2018; Parisot et al. 2018; Yang et al. 2019), our proposed graph neural network is novel in the following aspect. As opposed to these graph neural networks (Ktena et al. 2018; Parisot et al. 2018; Yang et al. 2019) that impose parameters on node features, our graph neural network directly imposes parameters on connectivity features instead. Imposing parameters on connectivity features is especially beneficial when the dimension of node features is very low, as it is the case that the node feature, i.e. the brain activation statistic, has only one dimension in our study. Hence, our graph neural network is well suitable for handling connectivity-driven problems, while the others mainly aim at dealing with node-driven problems.

The functional connectivity network has also been verified to transfer functional information across cortical regions (Cole et al. 2016; Ito et al. 2017). Under this activity flow mapping, functional activation information is transferred to neighboring brain regions via functional connectivity pathways. The activity flow mapping shares certain similarities with our study in the sense that the functional information propagates within the functional connectivity network. However, our study differs from the activity flow framework mainly in that functional activation information of all brain regions in our study is unknown, while only the ROI’s functional activation information is unknown in the activity flow framework. In this sense, our proposal and study require less functional information of brain regions and thus has practical implications in that one does not need to scan functional task contrasts beforehand to get the functional information of some brain regions.

Our results showed that the 2-layers graph neural network containing 1-hop and 2-hop functional connectivity best characterizes the rFFC region’s functional face response, indicating that 2-hops connectivity information may be enough to estimate the rFFC region’s function. On the other hand, from the computation perspective, as the number of layers in a graph neural network gets large, the parameters and complexity of the model also enlarge. Since the sample size is relatively limited compared to that of datasets in machine learning, models with large complexity are also likely to overfit the data and thus have a poor generalization ability. Future work involves utilizing datasets with a large sample size to test whether graph neural networks with more layers can further improve the generalization ability.

We chose the rFFC region that has a specialized function and is reliably replicated across studies to test our assumption primarily. However, the rFFC region is specialized in the face selectivity function which has special meaning in the human evolution process and has a specific neural mechanism (Tsao et al. 2006; Freiwald and Tsao 2014). Whether our proposal can be generalized to brain regions beyond the rFFC still remains to be solved, especially to brain regions that are more functionally variable across individuals and flexible across tasks, i.e. the heteromodal association cortices (Anderson et al. 2013; Mueller et al. 2013; Tei et al. 2017). Future work also includes extension to brain regions involving a wide functional domains to test our proposal.

We used undirected functional connectivity to construct the brain connectivity network in this study. However, the propagation of functional information in the brain is actually directional, and this directional information was not taken into account. Effective connectivity should be considered in the future to capture the directionality of information transfer. In addition, there are also other choices to construct the brain connectivity network, such as the structural connectivity representing white matter fiber pathways. Researchers can also explore the relationship between the multi-hops structural connectivity network and the individual brain region’s function.

In conclusion, we proposed that the multi-hops connectivity profile can improve the prediction performance of individual differences in the brain region’s function. Results revealed that the 2-hops functional connectivity network best characterizes the rFFC region’s individual functional face response. This advancement contributes to understanding the mechanism of individual brain region’s function in terms of the brain network and provides a new perspective on brain functional processing mechanisms at the network level.

## Acknowledgments

This work was supported by the National Key Research and Development Program of China (grant No. 2017YFB1002504). Data were provided by the Human Connectome Project, WU-Minn Consortium (Principal Investigators: David Van Essen and Kamil Ugurbil; 1U54MH091657) funded by the 16 NIH Institutes and Centers that support the NIH Blueprint for Neuroscience Research.

## Author contributions/Authorship

Wu, Li and Feng designed the research, Wu performed the data analysis, Wu, Li and Feng wrote and revised the paper.

