## Supplementary material for "Multi-hops functional connectivity improves individual prediction of fusiform face activation via a graph neural network"

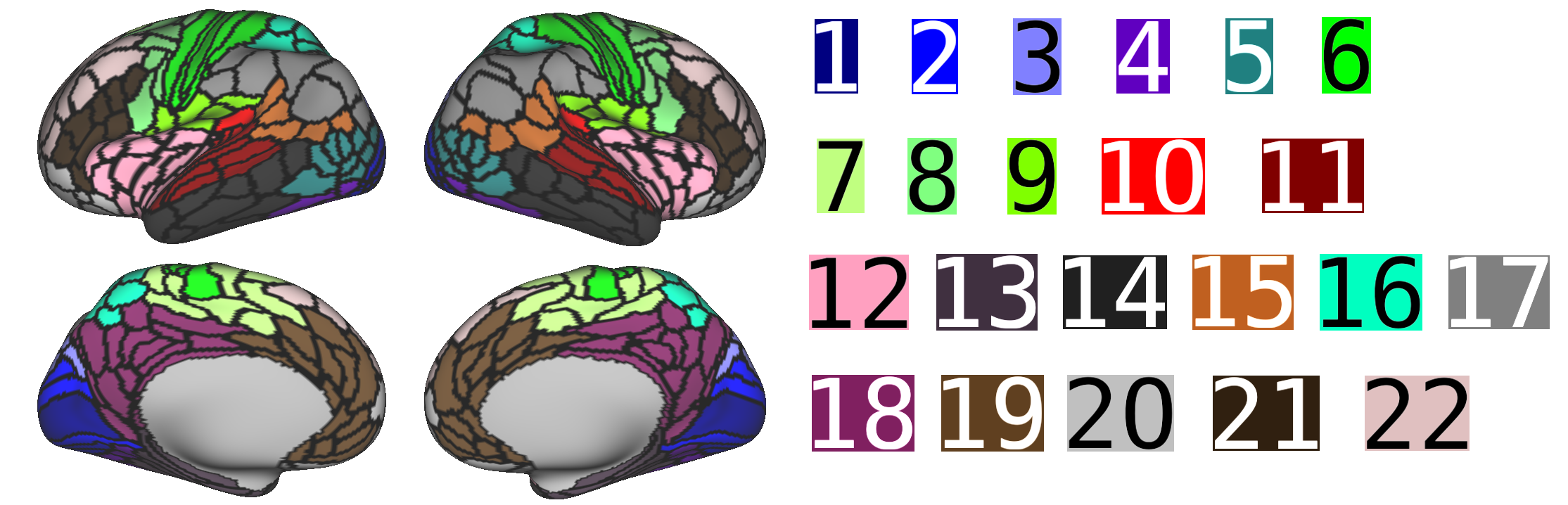

Figure S1. Color coding of the brain regions. The cortical names corresponding to the numbers are shown in Table S2. This figure is adapted from Figure 1 in the Supplementary Neuroanatomical Results for A Multi-modal Parcellation of Human Cerebral Cortex.

| FACES-SHAPES | | |
| --- | --- | --- |
| Mean | NSE | Correlation |
| Permutation | 1.032 | 0.011 |
| 1-GNN | 0.927 | 0.296 |
| 2-GNN | 0.857 | 0.392 |
| 3-GNN | 0.864 | 0.391 |
| Paired-sample *t*-test | NSE | Correlation |
| 2-GNN/Permutation | *t*(99) = -26.5,  *p* = 9.5×10^-47^ | *t*(99) = 33.2,  *p* = 1.7×10^-55^ |
| 2-GNN/1-GNN | *t*(99) = -16.0,  *p* = 3.0×10^-29^ | *t*(99) = 13.5,  *p* = 2.8×10^-24^ |
| 2-GNN/3-GNN | *t*(99) = -2.09,  *p* = 0.039 | *t*(99) = 0.269,  *p* = 0.788 |
| 3-GNN/1-GNN | *t*(99) = -10.4,  *p* = 1.6×10^-17^ | *t*(99) = 11.5,  *p* = 4.7×10^-20^ |
| FACE-AVG | | |
| Mean | NSE | Correlation |
| Permutation | 1.024 | 0.008 |
| 1-GNN | 0.933 | 0.291 |
| 2-GNN | 0.886 | 0.355 |
| 3-GNN | 0.915 | 0.332 |
| Paired-sample *t*-test | NSE | Correlation |
| 2-GNN/Permutation | *t*(99) = -20.9,  *p* = 3.9×10^-38^ | *t*(99) = 25.3,  *p* = 4.5×10^-45^ |
| 2-GNN/1-GNN | *t*(99) = -11.3,  *p* = 1.4×10^-19^ | *t*(99) = 9.29,  *p* = 3.8×10^-15^ |
| 2-GNN/3-GNN | *t*(99) = -7.93,  *p* = 3.4×10^-12^ | *t*(99) = 5.96,  *p* = 3.9×10^-8^ |
| 3-GNN/1-GNN | *t*(99) = -2.70,  *p* = 0.008 | *t*(99) = 4.65,  *p* = 1.0×10^-5^ |

Table S1. Statistical comparison of the prediction metrics. The mean prediction metrics and the statistical comparison of metrics between different models were shown. Permutation represents the random permutation model. The k-GNN represents the graph neural network with k layers. NSE is the abbreviation of normalized squared error.

| Number of cortex | Full name | Abbreviation |
| --- | --- | --- |
| 1 | Primary_Visual | Primary_V |
| 2 | Early_Visual | Early_V |
| 3 | Dorsal_Stream_Visual | Dorsal_Stream_V |
| 4 | Ventral_Stream_Visual | Ventral_Stream_V |
| 5 | MT+_Complex_and_Neighboring_Visual_Areas | MT+ |
| 6 | Somatosensory_and_Motor | - |
| 7 | Paracentral_Lobular_and_Mid_Cingulate | - |
| 8 | Premotor | Premotor |
| 9 | Posterior_Opercular | - |
| 10 | Early_Auditory | - |
| 11 | Auditory_Association | - |
| 12 | Insular_and_Frontal_Opercular | - |
| 13 | Medial_Temporal | Medial_T |
| 14 | Lateral_Temporal | Lateral_T |
| 15 | Temporo-Parieto-Occipital_Junction | TPO_J |
| 16 | Superior_Parietal | Superior_P |
| 17 | Inferior_Parietal | Inferior_P |
| 18 | Posterior_Cingulate | Posterior_C |
| 19 | Anterior_Cingulate_and_Medial_Prefrontal | - |
| 20 | Orbital_and_Polar_Frontal | OP_F |
| 21 | Inferior_Frontal | Inferior_F |
| 22 | Dorsolateral_Prefrontal | Dorsolateral_PF |

Table S2. Full names of the abbreviations used in Figure 4. Only the cortices that appeared in the article are abbreviated. These names were used in the Supplementary Neuroanatomical Results for A Multi-modal Parcellation of Human Cerebral Cortex.
